# Differences in Long-Chained Cuticular Hydrocarbons between Males and Gynes in *Cataglyphis* Desert Ants

**DOI:** 10.1101/538926

**Authors:** Shani Inbar, Eyal Privman

## Abstract

Cuticular hydrocarbons play an important role in chemical communication in social insects, serving, among other things, as nestmate, gender, dominance and fertility recognition cues. In ants, however, very little is known about the precopulatory signals cuticular hydrocarbons carry. These signals may serve as affecting sex pheromones and aphrodisiacs or as reliable signals for idiosyncratic traits, which indirectly affect sexual selection. In this study, we examined, for the first time in the *Cataglyphis* genus, sex-specific variability in cuticular hydrocarbons. We focused on a population in which we observed either unmated queens or males, but never both, present in the same colony before mating season. We found significant quantitative differences, but no qualitative differences, between virgin queens and their potential mates. In analyses of both absolute amounts and relative amounts of cuticular hydrocarbons, we found different compounds to be significantly displayed on gynes and drones, suggesting absolute and relative amounts may carry different signals influencing mating behavior and mate choice. We discuss the possible signals advertised by the non-polar fraction of these hydrocarbon profiles.

## INTRODUCTION

While in most animals sexual selection presents itself in many elaborate and colorful ways, in social insects, pre-mating sexual selection is considered to be limited (Boomsma *et al.* 2005). And still, there are various mechanisms through which sexual selection acts that have been demonstrated in social insects, and in ants specifically, most of which are post-mating, such as sperm selection, use of mating plugs, and increased mating frequencies by both males and females (see overview in Boomsma *et al.* (2005)). Precopulatory selection in ants, however, is limited to few cases of male territoriality (Abell *et al.* 1999; Davidson 1982; Heinze *et al.* 1998; Kinomura and Yamauchi 1987; Stuart *et al.* 1987; Wiernasz *et al.* 2001) and semiochemical communication, which has been surprisingly understudied (Ayasse *et al.* 2001).

Some studies have shown virgin queens to be advertising sexual receptivity (Reviewed in Ayasse *et al.* (2001) and Hölldobler and Wilson (1990)) and others have identified active female sex pheromones in several species (*Formica lugubris* in Walter *et al.* (1993), *Polyergus breviceps* in Greenberg *et al.* (2007) and in Greenberg *et al.* (2018), *Polyergus rufescens* in Castracani *et al.* (2005) and in Castracani *et al.* (2008)). Mandibular, Dufour, poison, and pygidial glands have also been shown to contain attractants to drones (Grasso *et al.* 2003). In several species, drones have been shown to discharge their mandibular glands during nuptial flights or when leaving the nest (BENTO *et al.* 2007; Brand *et al.* 1973b; Hölldobler 1976; Law *et al.* 1965) and male metapleural glands were also suggested to have an active role in sexual selection (Hölldobler and Engel-Siegel 1984).

Cuticular hydrocarbons (CHCs) are likely to play an important role in mating behavior and mate choice in ants because many other individual and colonial difference such as task (see for example Greene and Gordon (2003)), caste and sex (see for example Campos *et al.* (2012)), nestmate recognition (See for example Lahav *et al.* (1999)) and fertility (See for example Dietemann *et al.* (2003)) are advertised by the cuticular chemical coating. Part of the chemical bouquet displayed on the cuticle may be active sex pheromones, but it can also contain signals indicative of other traits which influence mating behavior. In ants, few studies demonstrated unique sex-specific differences in CHCs between virgin queens (hereafter gynes) and males (drones). Those studies showed these differences to be either qualitative or quantitative (Antonialli Junior *et al.* 2007; Brand *et al.* 1973a; Chernenko *et al.* 2012; Cremer *et al.* 2002; Cuvillier-Hot *et al.* 2001; Hojo *et al.* 2008; Johnson and Sundström 2012; Kureck *et al.* 2011; Oppelt *et al.* 2008).

To the best of our knowledge, no study examined sexual dimorphism in CHCs in *Cataglyphis* ants and the mechanisms underlying sexual selection are still unknown in this large genus. In *C. iberica* a mixture of three linear alkanes and methylalkanes (n-C27, nC29 and 3-meC29) has been suggested to function as queen pheromones but their function was restricted to sterility inducement rather than male attraction (Van Oystaeyen *et al.* 2014). Another study found that gynes produce more undecane in the Dufour’s gland than drones (Monnin *et al.* 2018) but also suggested it may have functions unrelated to male attraction.

Another possible mechanism related to precopulatory sexual selection may involve altering acceptance threshold making drones able to get closer to gynes without being harassed by workers (Helft *et al.* 2016; Helft *et al.* 2015). There is, indeed, evidence of drones mimicking the queen’s chemical bouquet in ants, thus allowing them to escape aggression from other drones (Cremer *et al.* 2002) or from workers (Franks and Hölldobler 1987). Other such acceptance-altering effects have been demonstrated in honeybees where different nestmate recognition mechanisms are used to identify drones and workers (Moritz and Neumann 2004), enabling drones to immigrate to foreign colonies.

Kinship between gynes and drones may also influence mate choice. Although it has been argued that discriminatory abilities are usually limited to nestmate recognition, regardless of kinship degree (Carlin 1988; Grafen 1990), there is evidence of the influence of kinship on mating preferences in ants (Keller and Passera 1993). Keller and Passera (1993), in this study, suggested that colony-derived cues may be of less importance in mating preferences than kinship cues. In bees, male have also been shown to discriminate between female kin through olfactory signals, such as CHCs (Smith 1983) suggesting that kinship recognition is, indeed, of importance in mate choice.

CHC may carry other signals which affect pre-mating sexual selection in ants such as overall fitness of both gynes and drones. Body size, in both sexes, is a reliable indicator of fitness and, in *Pogonomyrmex* harvester ants, for example, larger drones have been documented to be more successful in mating attempts (Abell *et al.* 1999; Wiernasz *et al.* 2001; Wiernasz *et al.* 1995). In ants, gynes and drones vary significantly in body size, ranging from both sexes being about equal in size to queens having ∼3 times the body length and ∼25 times the body mass of males (Boomsma *et al.* 2005). Cataglyphis ants show significant variability in body size between gynes and drones.

Body size and surface area effect the absolute amounts of CHCs coating an individual ant and the total amount of a component is proportional to it. It is hard to determine whether a factor in the pheromonal blend is informative because of its absolute amount, its ratio with other chemicals, or both factors combined. Ratios between CHCs are usually less variable than absolute quantities of each compound (See for example in Drosophila Coyne (1996)) because absolute amounts are affected by body size. The opposite case was also reported, with higher variability in relative amounts than in total amounts (See for example, also in Drosophila, Grillet *et al.* (2005)). Thus, body size variation of both gynes and drones may play a role in mating. We thus chose a twofold analysis of the CHCs to account for the possibility of absolute amounts enhancing differences between gynes and drones in ants of *C. niger* and also the possibility of them masking differences. To our knowledge, this is the first study to demonstrate sexual dimorphism in CHCs in *Cataglyphis* and it suggests that ratios between compounds may carry different signals than absolute amounts.

## METHODS AND MATERIALS

### Ants

Ants were collected between April 24 and May 10, 2016 from colonies dug along a 4 km transect in Betzet beach on the northern Israeli coastline (from N33.05162, E35.10245 to N33.07868, E35.10705). This population was previously referred to as *Cataglyphis drusus* (Eyer *et al.* 2017) but our recent species delimitation study suggested that this is the same species as *C. niger*, because these populations are not differentiated by their nuclear genomic DNA (Brodetzki *et al.* 2019). Colonies of this population are monogyne (headed by a single queen) and polyandrous (queens are multiply mated) (Brodetzki *et al.* 2019; Eyer *et al.* 2017). Queens usually mate during the spring and sexuals (gynes and drones) can usually be found in nests in April-May. In this study, on the days when ants were collected, we observed colonies with either gynes or drones (or neither of the two) present in the colony before mating season. We used samples from twelve nests, 6 female-producing colonies and 6 male-producing colonies, with 1-3 gynes/drones collected from each colony. A total of 13 gynes and 10 drones were analyzed. All sexuals were frozen on the same evening of collection.

### Cuticular hydrocarbon analysis

Whole bodies were individually immersed in hexane, containing 400 ng/µl of tetracosane (C24) as internal standard for quantitative analysis. Initial analysis was conducted by gas chromatography/mass spectrometry (GC/MS), using a VF-5ms capillary column, temperature-programmed from 60°C to 300°C (with 1 min initial hold) at a rate of 10°C per min, with a final hold of 15 min. Compound were identified according to their fragmentation pattern and respective retention time compared to authentic standards. We identified 34 compounds in gynes and the same 34 compounds in drones. Quantitative analyses for each individual were performed by flame ionization gas chromatography (GC/FID), using the above running conditions. Peak integration was performed using the program Galaxie Varian 1.9.

### Data and Statistical analysis

Statistical analysis was performed using XLSTAT (https://www.xlstat.com; Addinsoft 2019;Boston, USA).

## RESULTS

Chemical analysis of the non-polar fraction of the cuticular extracts of gynes and drones identified 34 long-chained CHCs, ranging from pentacosane (c25) to tritriacontane (c33). All 34 compounds were identified in both gynes and drones with no qualitative differences between them (Fig. 1). Quantitative differences between the two groups were examined in two ways: first, an analysis of absolute quantities calculated relative to an internal standard; second, an analysis of the percentage of every compound in the total extract, calculated according to percent area in peak integration (Table 1). Principal component analyses (PCA) of both measures are shown in Figure 2. Colony level variation did not seem to mask the differences between gynes and drones’ cuticular profiles as shown in Figure 2.

**Table 1:** Quantification of CHCs of 13 gynes (from 6 colonies) and 10 drones (from 6 colonies)

|  |  | Gynes<br>Absolute quantity<br>(ng) | Drones<br>Absolute quantity<br>(ng) | P- Value | Gynes<br>% of total<br>secretion | Drones<br>% of total<br>secretion | P- Value |
| --- | --- | --- | --- | --- | --- | --- | --- |
| 1 | c25 | 1169.10±780.33<br>(239.73 - 3288.89) | 865.49±1345.39<br>(78.36 - 4512.50) |  | 1.80±0.66<br>(1.14 - 3.45) | 2.74±0.93<br>(1.37 - 3.96) | 0.010 |
| 2 | 5me c25 | 108.92±104.41<br>(24.66 - 400.00) | 69.43±69.84<br>(1.51 - 237.50) | 0.046 | 0.15±0.06<br>(0.07 - 0.27) | 0.38±0.42<br>(0.01 - 1.40) |  |
| 3 | 3me c25 | 4040.73±4316.35<br>(528.77 - 17266.67) | 1083.44±879.80<br>(181.29 - 3112.50) | 0.006 | 5.27±1.65<br>(3.37 - 8.14) | 5.17±2.73<br>(2.49 - 9.12) |  |
| 4 | c26 | 768.13±363.13<br>(363.01 - 1577.78) | 311.59±333.96<br>(31.83 - 912.94) |  | 1.35±0.59<br>(0.71 - 2.67) | 1.21±0.59<br>(0.49 - 2.23) |  |
| 5 | 10me c26 | 286.50±373.59<br>(17.81 - 1422.22) | 35.19±33.87<br>(5.26 - 115.94) | 0.024 | 0.33±0.18<br>(0.10 - 0.64) | 0.22±0.18<br>(0.02 - 0.46) |  |
| 6 | 4me c26 | 401.26±408.87<br>(67.12 - 1622.22) | 83.34±61.54<br>(9.00 - 187.50) |  | 0.53±0.13<br>(0.35 - 0.73) | 0.42±0.26<br>(0.14 - 0.86) |  |
| 7 | 4,12 dime c26 | 171.95±161.97<br>(20.55 - 622.22) | 44.34±38.63<br>(7.61 - 137.50) |  | 0.22±0.06<br>(0.15 - 0.30) | 0.20±0.10<br>(0.11 - 0.43) |  |
| 8 | c27 | 822.32±408.46<br>(298.63 - 1844.44) | 1170.72±2631.94<br>(84.43 - 8587.50) |  | 1.38±0.43<br>(0.68 - 2.20) | 2.33±1.70<br>(0.79 - 6.89) |  |
| 9 | 11+13 me c27 | 6150.91±6536.77<br>(702.74 - 25200.00) | 1425.94±1218.75<br>(147.40 - 4100.00) | 0.038 | 7.72±2.67<br>(4.72 - 11.36) | 6.49±3.42<br>(2.27 - 11.21) |  |
| 10 | 5me c27 | 70.74±56.68<br>(11.94 - 222.22) | 26.88±30.06<br>(0.92 - 100.00) |  | 0.12±0.08<br>(0.02 - 0.28) | 0.12±0.08<br>(0.01 - 0.26) |  |
| 11 | 11,15 dime c27 | 2254.88±3164.19<br>(95.89 - 11311.11) | 373.00±415.91<br>(3.46 - 1405.80) | 0.011 | 2.44±2.09<br>(0.69 - 6.54) | 2.00±2.16<br>(0.06 - 5.32) |  |
| 12 | 7,11 dime c27 | 475.86±448.06<br>(60.27 - 1785.71) | 73.76±65.60<br>(4.84 - 220.38) |  | 0.72±0.41<br>(0.15 - 1.51) | 0.44±0.42<br>(0.07 - 1.50) |  |
| 13 | 3me c27 | 2017.75±1746.58<br>(445.21 - 7133.33) | 924.67±1204.11<br>(131.58 - 4137.50) | 0.004 | 2.85±0.55<br>(1.85 - 3.39) | 3.31±0.66<br>(2.42 - 4.43) |  |
| 14 | 7,11,15 trime c27 | 675.95±521.63<br>(83.56 - 1888.89) | 124.08±100.80<br>(6.23 - 312.50) | 0.021 | 0.93±0.18<br>(0.62 - 1.32) | 0.63±0.36<br>(0.10 - 1.02) | 0.016 |
| 15 | c28 | 1327.92±956.33<br>(301.37 - 3666.67) | 470.54±589.69<br>(71.97 - 2062.50) | 0.011 | 1.93±0.21<br>(1.65 - 2.36) | 1.76±0.39<br>(1.11 - 2.47) |  |
| 16 | 12me c28 | 1652.54±1337.81<br>(272.60 - 5044.44) | 421.62±395.33<br>(59.52 - 1387.50) |  | 2.27±0.34<br>(1.81 - 3.14) | 1.83±0.65<br>(0.92 - 2.94) |  |
| 17 | 4me+ 8,12 dime c28 | 1701.79±1448.09<br>(304.11 - 5666.67) | 393.80±344.16<br>(56.75 - 1187.50) |  | 2.31±0.18<br>(2.01 - 2.65) | 1.79±0.65<br>(0.87 - 2.58) |  |
| 18 | 2me c28 | 60.12±45.04<br>(14.93 - 155.56) | 25.03±33.88<br>(3.46 - 100.00) |  | 0.09±0.04<br>(0.04 - 0.15) | 0.08±0.03<br>(0.05 - 0.16) |  |
| 19 | 4,12 dime c28 | 200.57±162.33<br>(32.88 - 577.78) | 32.42±25.33<br>(9.69 - 98.55) |  | 0.28±0.04<br>(0.22 - 0.35) | 0.20±0.11<br>(0.02 - 0.34) |  |
| 20 | c29 | 1350.87±663.95<br>(334.38 - 2444.44) | 3601.76±8320.86<br>(155.76 - 26875.00) |  | 2.36±1.16<br>(0.90 - 4.50) | 6.09±6.17<br>(1.21 - 21.57) |  |
| 21 | 11+13 me c29 | 12893.58±9321.57<br>(2461.64 - 35444.44) | 5024.77±6563.04<br>(803.51 - 22350.00) | 0.014 | 18.38±1.78<br>(15.05 - 21.01) | 17.31±1.98<br>(14.32 - 20.82) |  |
| 22 | 11,15 dime c29 | 8757.70±6757.88<br>(66.67 - 26628.57) | 2583.85±2871.68<br>(532.75 - 10000.00) |  | 14.38±5.04<br>(0.03 - 19.07) | 10.30±2.02<br>(7.21 - 14.51) | 0.025 |
| 23 | 3me c29 | 4347.20±10298.52<br>(575.00 - 38511.11) | 864.98±981.86<br>(182.46 - 3000.00) | 0.014 | 3.86±4.14<br>(1.57 - 17.36) | 3.26±0.78<br>(2.30 - 4.96) |  |
| 24 | 7,11,15 trime c29 | 96.00±86.39<br>(6.25 - 314.29) | 20.99±18.17<br>(4.61 - 66.67) | 0.025 | 0.13±0.06<br>(0.02 - 0.22) | 0.12±0.07<br>(0.02 - 0.23) |  |
| 25 | c30 | 2255.41±1856.09<br>(494.52 - 6977.78) | 712.55±873.58<br>(178.55 - 3050.00) | 0.018 | 3.10±0.46<br>(2.04 - 3.78) | 2.73±0.40<br>(2.15 - 3.38) |  |
| 26 | 14 me c30 | 1755.93±1438.75<br>(287.67 - 5355.56) | 514.17±593.81<br>(141.94 - 2075.00) |  | 2.37±0.25<br>(1.93 - 2.67) | 2.06±0.48<br>(1.46 - 2.72) |  |
| 27 | 4 me c30 | 1773.93±1566.14<br>(353.42 - 5911.11) | 432.60±407.25<br>(145.62 - 1437.50) | 0.017 | 2.37±0.36<br>(1.62 - 2.79) | 1.95±0.52<br>(1.15 - 2.70) |  |
| 28 | 2 me c30 | 311.34±321.04<br>(63.01 - 1177.78) | 45.75±33.89<br>(18.43 - 113.04) |  | 0.40±0.10<br>(0.21 - 0.56) | 0.26±0.13<br>(0.07 - 0.45) | 0.011 |
| 29 | c31 | 276.17±165.32<br>(31.25 - 566.15) | 562.24±1278.31<br>(19.35 - 4150.00) |  | 0.51±0.36<br>(0.10 - 1.17) | 1.14±0.91<br>(0.16 - 3.33) | 0.035 |
| 30 | 13+15 me c31 | 3413.97±2341.85<br>(709.59 - 8400.00) | 2003.26±3378.76<br>(251.61 - 11262.50) | 0.019 | 4.96±1.15<br>(3.05 - 6.66) | 6.12±2.88<br>(2.85 - 10.66) |  |
| 31 | 11,15 dime c31 | 9198.67±6728.53<br>(2210.96 - 24933.33) | 3385.37±3031.83<br>(985.26 - 9477.65) |  | 13.32±3.24<br>(7.79 - 18.57) | 16.09±7.76<br>(6.15 - 30.50) |  |
| 32 | 7,11,15 trime c31 | 439.66±366.76<br>(65.63 - 1333.33) | 95.61±106.64<br>(18.43 - 375.00) |  | 0.62±0.20<br>(0.21 - 0.89) | 0.50±0.33<br>(0.07 - 1.02) |  |
| 33 | c32 | 392.66±421.94<br>(41.18 - 1400.00) | 133.82±114.07<br>(23.96 - 375.00) |  | 0.49±0.24<br>(0.07 - 0.86) | 0.69±0.49<br>(0.28 - 1.75) |  |
| 34 | c33 | 8.60±11.22<br>(0.00 - 42.86) | 20.65±34.13<br>(0.00 - 112.50) |  | 0.05±0.11<br>(0.00 - 0.41) | 0.08±0.11<br>(0.00 - 0.37) |  |

Means ± Standard deviation are shown and ranges are given in parentheses.

**Figure 1:**
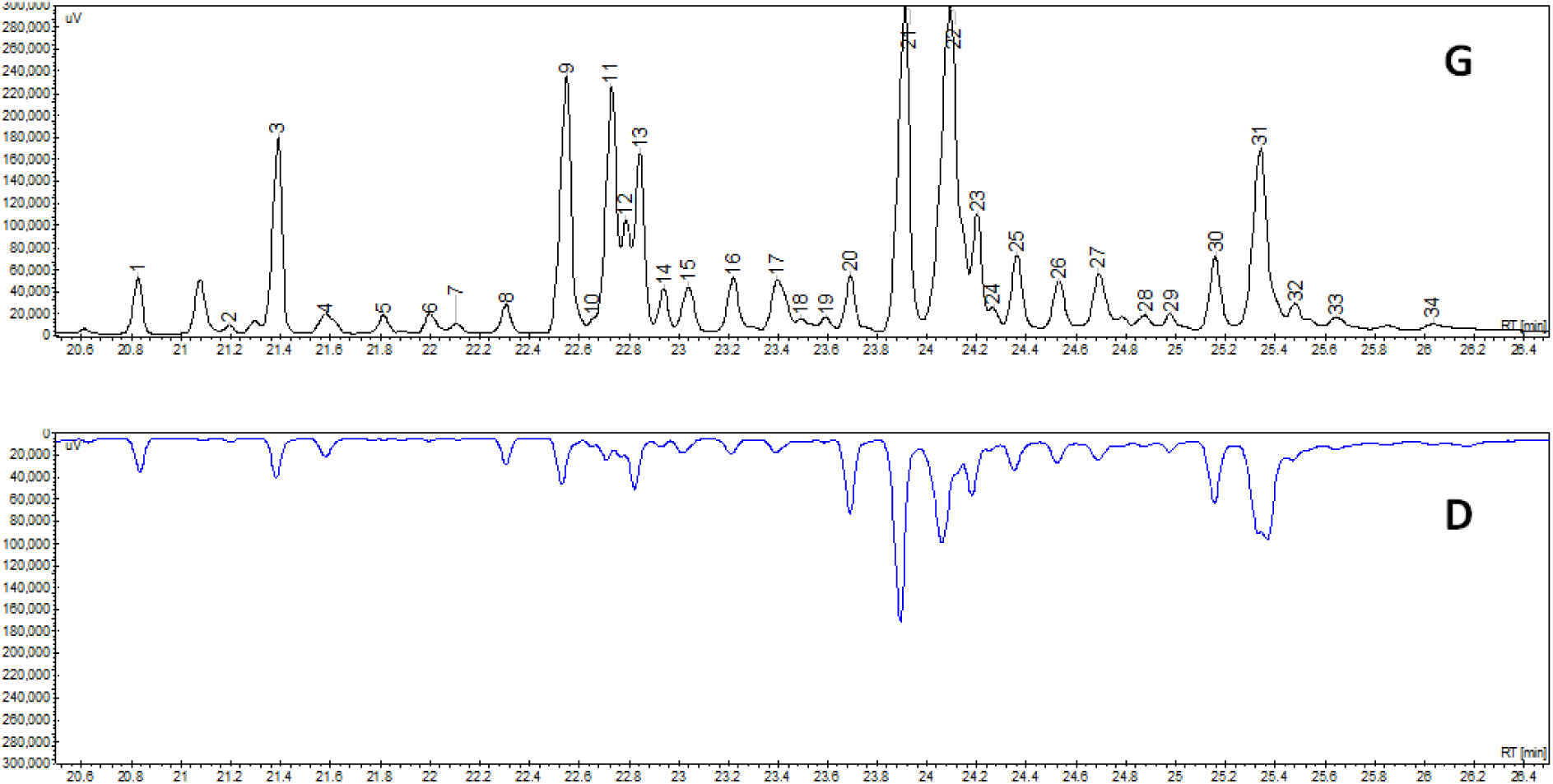
Chromatograms of CHCs from total body extracts. The upper chromatogram is of a gyne and the lower of a drone. Only long-chained CHC are shown.

**Figure 2:**
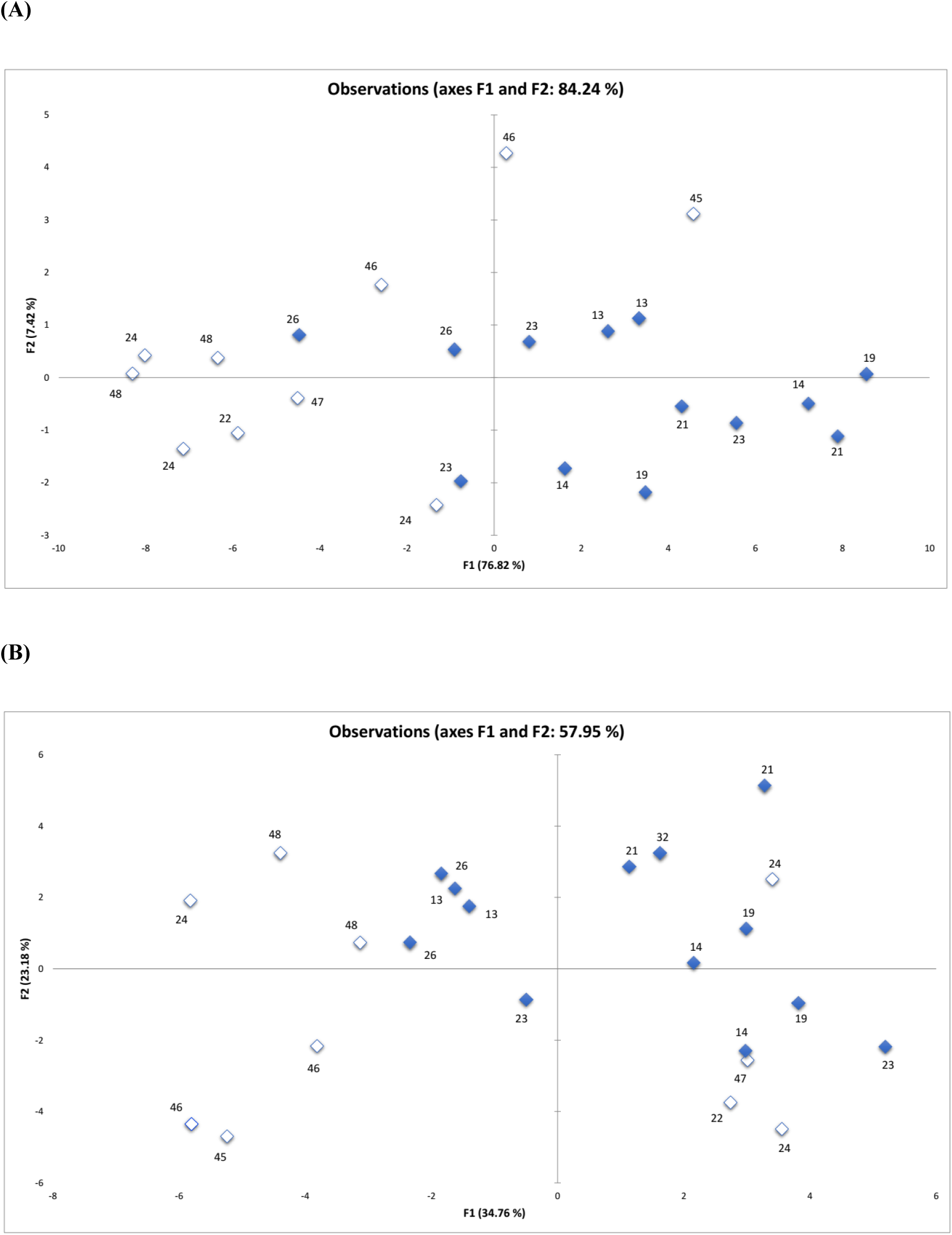
Principal component analysis (PCA). **(A)** Absolute quantities **(B)** Percent of total extract. Filled blue points represent individual observations of gynes and empty points of drones. Numbers indicate colony identity.

In a linear discriminant analysis (LDA), both relative amounts and absolute quantities showed significant differences between CHC profiles of gynes and drones (In Wilks’ Lambda test (Rao’s approximation) *p*-values were 0.044 and 0.002, for absolute amounts and relative amounts respectively). Relative amounts were less variable than absolute amounts. Following an LDA analysis, a unidimensional test of equality of the means of the groups showed different compounds to be significantly discriminating between gynes and drones in the two analyses; four compounds in the relative amount analysis: C25; 11,15-dime C29; 2-me C30; C31; and 13 compounds in the absolute amounts analysis: 5-me C25; 3-me C25; 10-me C26; 11+13-me C27; 11,15-dime C27; 3-me C27; C28; 11+13-me C29; 3-me C29; 7,11,15-trime C29; C30; 4-me C30; 13+15-me C31. Only one compound was identified in both analyses - 7,11,15-trime C27 (Table 1).

## DISCUSSION

Our study reveals differences in long-chained CHCs between drones and gynes in *Cataglyphis niger* colonies. The differences were quantitative and not qualitative and different compounds stand out in a discriminant analysis when considering either relative amounts or absolute amounts. The two analyses revealed different compounds to be significantly discriminating virgin queens and their potential mates.

We reported that on the days when ants were collected, we observed colonies with either gynes or drones (or neither of the two) which may suggest that this species exhibit a split sex ratio (Boomsma 1991; Boomsma and Grafen 1990, 1991; Bourke 2005; Chan and Bourke 1994; Chapuisat and Keller 1999; Grafen 1986; Mehdiabadi *et al.* 2003; Pamilo and Seppä 1994; Queller and Strassmann 1998; Ratnieks and Boomsma 1997; Sundström 1994). As (Trivers and Hare 1976) suggested, split sex-ratio may be the result of a tug-of-war between queens and workers over reproductive control. There is reason to believe that this is the case in our population because previous studies suggest that in colonies with singly-mated queens, queens favor a 1:1 ratio of gynes to drones and workers a 3:1 ratio. Although studies from singly-mated species revealed frequent female-biased sex ratios indicating that workers often prevail (Herbers 1984; Nonacs 1986; Trivers and Hare 1976), the queen may thwart worker control by specializing in producing either gynes or drones. However, determining whether colonies exhibit split sex-ratios requires a long-term study. It is possible that colonies alternate between female and male production in different years or that the production is temporally offset, so that drones are produced slightly before gynes or vice versa (Keller *et al.* 1996). Colonies, therefore, should be thoroughly examined at different times in the reproductive season and in multiple years. We have not done so in the present study because our goal was different: to determine differences in circular hydrocarbons between gynes and their potential mates as a potential mechanism involved in sexual selection.

We reported using 1-3 gynes or drones from each of 6 female and 6 male producing colonies. This sample size may seem small in light of the number of variables (34 compounds) that were identifies, however because CHC’s are produced in series (Martin and Drijfhout 2009), the number of variables is, in fact, smaller and a large sample size is not necessary in order to determine statistical significance.

Interestingly, when results were compared to previous studies in ants, similar discriminatory compounds could be found. The three linear alkanes and methylalkanes found in *C. iberica* (Van Oystaeyen *et al.* 2014) were also present in *C. niger* but only 3-meC29 was statistically different between gynes and drones in both species. This difference was only found in our absolute amounts analysis and not in the relative amounts analysis. Six CHC’s which were identified in our analysis were also identified in *Formica fusca* (Chernenko *et al.* 2012) (3-MeC25, 4-MeC26, 11,15-diMeC27, 3-MeC27 12-MeC28, 8,12-diMeC28, C29) but only three of them were statistically different between gynes and drones in both species (3-MeC25, 11,15-diMeC27, 3-MeC27). Here again, these difference were only found in our absolute amounts analysis and not in our relative amounts analysis. We interpret these findings as an indication of absolute amount carrying additional information involved in sexual selection, while not masking information carried by relative amounts.

Two main issues can be addressed in light of our results:

First, although bioassays are needed in order to demonstrate active sex pheromones, the significant differences between gynes and drones suggest that CHCs carry signals related to mating. These chemical cues may play different roles such as altering acceptance threshold, acting as reliable signals of fitness and fertility or kinship. Our results also show that there is an abundance of branched hydrocarbons in the CHC coating of both gynes and drones and it is known that such compounds reduce the waterproofing efficacy of the cuticle (Gibbs and Pomonis 1995). This reduced efficiency may, therefore, also play a role in signaling fitness, through the handicap principle (Heinze and d’Ettorre 2009; Zahavi and Zahavi 1999) as it has been suggested before (Boulay *et al.* 2017).

Second, we have shown that it is important to analyze both total amounts and relative amounts. Our study showed them both to be significantly different between gynes and drones. Absolute and relative amounts may carry distinct cues; Absolute quantities may be indicative of body size, carrying signals of overall fitness and fertility, while relative amount may be cues for kinship. Therefore, it is not unlikely that both absolute quantities and relative amounts influence mate choice and mating behavior and that both drones and gynes integrate multiple cues. We encourage future studies to account for both factors and hope that future bio-assays will help identify specific sex pheromones as well as other signals affecting sexual behavior and mating in the *Cataglyphis* genus.

## Acknowledgement

We thank Abraham Hefetz for introducing us to these wonderful ants and for assistance with the chemical analysis. E.P. was supported by Israel Science Foundation Grants no. 646/15, 2140/15, and 2155/15 and US-Israel Binational Science Foundation Grant no. 2013408

